# SV-plaudit: A cloud-based framework for manually curating thousands of structural variants

**DOI:** 10.1101/265058

**Authors:** Jonathan R. Belyeu, Thomas J. Nicholas, Brent S. Pedersen, Thomas A. Sasani, James M. Havrilla, Stephanie N. Kravitz, Megan E. Conway, Brian K. Lohman, Aaron R. Quinlan, Ryan M. Layer

**Affiliations:** Department of Human Genetics, University of Utah, Salt Lake City, UT; USTAR Center for Genetic Discovery, University of Utah, Salt Lake City, UT; Department of Biomedical Informatics, University of Utah, Salt Lake City, UT; To whom correspondence should be addressed

**Keywords:** Structural variants, Visualization, Manual curation

## Abstract

*SV-plaudit* is a framework for rapidly curating structural variant (SVs) predictions. For each SV, we generate an image that visualizes the coverage and alignment signals from a set of samples. Images are uploaded to our cloud framework where users assess the quality of each image using a client-side web application. Reports can then be generated as a tab-delimited file or annotated VCF. As a proof of principle, nine researchers collaborated for one hour to evaluate 1,350 SVs each. We anticipate that *SV-plaudit* will become a standard step in variant calling pipelines and the crowd-sourced curation of other biological results.

Code available at https://github.com/jbelyeu/SV-plaudit

Demonstration video available at https://www.youtube.com/watch?v=ono8kHMKxDs

## INTRODUCTION

Large genomic rearrangements, or structural variants (SVs), are an abundant form of genetic Variation within the human genome^2^, and they play an important role in both species evolution^3,4^ and human disease phenotypes^5-9^. While many methods have been developed to identify SVs from whole-genome sequencing (WGS) data^10-14^, the accuracy of SV prediction remains far below that of single-nucleotide and insertion-deletion variants^1^. Improvements to SV detection algorithms have, in part, been limited by the availability and applicability of high-quality truth sets. While the Genome in a Bottle^15^ consortium has made considerable progress toward a goldstandard variant truth set, the incredibly high quality of the data underlying this project (300X and PCR-free) calls into question the generality of the accuracy they obtain in typical quality WGS datasets (30X with PCR-amplification).

Given the high false positive rate of SV calls from genome and exome sequencing, manual inspection is a critical quality control step, especially in clinical cases. Scrutiny of the evidence supporting an SV is considered to be a reliable "dry bench" validation technique, as the human eye can rapidly distinguish true SV signal from alignment artifacts. In principle, we could improve the accuracy of SV call sets by visually validating every variant. In practice, however, current genomic data visualization methods^16-20^ were designed primarily for spot checking a small number of variants and are difficult to scale to the thousands of SVs in typical call sets. Therefore, a curated set of SVs requires a new framework that scales to thousands of SVs, minimizes the time needed to adjudicate individual variants, and manages the collective judgment of large and often geographically dispersed teams.

Here we present *SV-plaudit,* a fast, highly-scalable framework enabling teams of any size to collaborate on the rapid, web-based curation of thousands of SVs. While support for teams of curators is not strictly required, collecting multiple opinions for each SV allows *SV-plaudit* to report the consensus view (i.e., a "curation score") of each variant. This consensus is less susceptible to human error and does not require expert users to score variants. With *SV-plaudit,* it is practical to inspect and score every variant in a call set, thereby improving the accuracy of SV predictions in individual genomes, and curating high quality-truth sets for SV method tuning.

## METHODS

### Overview

*SV-plaudit* (Fig 1A) is based on two software packages: ***samplot*** for SV image generation, and ***PlotCritic*** for staging the Amazon cloud environment and managing user input. Once the environment is staged, users log into the system and are presented with a series of SV images in either a random or predetermined order. For each image, the user answers the curation question and responses are logged. Reports on the progress of a project can be quickly generated at any point in the process.

**Figure 1.**
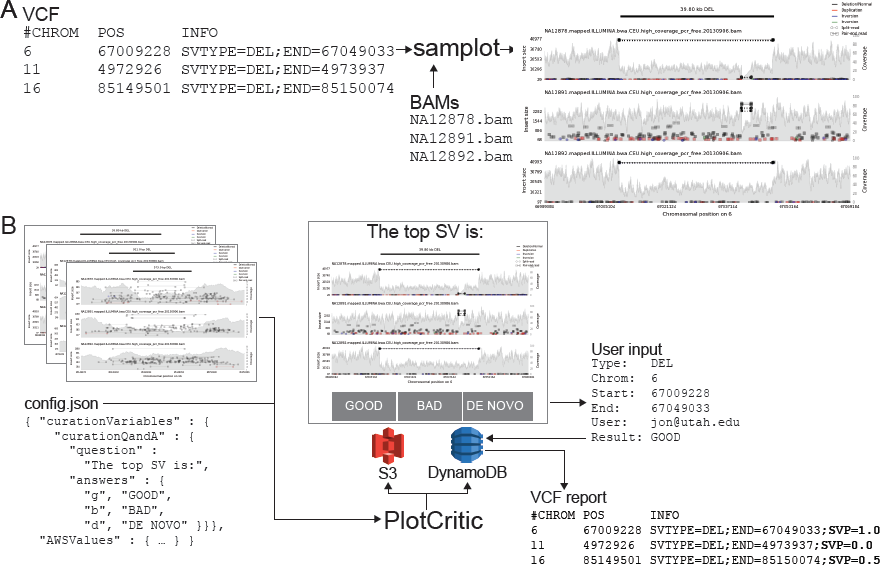
The *SV-Plaudit* process. **A)** *Samplot* generates an image for each SV from VCF considering a set of alignment (BAM or CRAM) files. **B)** *PlotCritic* uploads the images to an Amazon S3 bucket and prepares DynamoDB tables. Users select a curation question (“GOOD”, "BAD”, or “DE NOVO”) for each SV image. DynamoDB logs user responses and generates reports. Within a report, a curation score function can be specified by mapping answer options to values and selecting an aggregation function. Here “GOOD” and “DE NOVO” were mapped to one, “BAD” to zero, and the mean was used. One useful output option for a report is a VCF annotated with the curation scores (shown here in bold as a **SVP**).

### Samplot

Samplot is a Python program that uses *pysam^21^* to extract alignment data from a set of BAM or CRAM files, and *matplotlib^22^* to visualize the raw data for the genomic region surrounding a candidate SV (Fig 1A). For each alignment file, *samplot* renders the depth of sequencing coverage, paired-end alignments, and split-read alignments where paired-end and split-read alignments are color-coded based by the type of SV they support (e.g., black for deletion, red for a duplication, etc.) (Fig 2). Alignments are positioned along the x-axis by genomic location and along the left y-axis by the distance between the ends (insert size), which helps users to differentiate normal alignments from discordant alignments that support an SV. Depth of sequencing coverage is also displayed on the right y-axis to allow users to inspect whether putative copy number changes are supported by the expected changes in coverage. To improve performance for large events, we downsample “normal” paired-end alignments (a +/− orientation and an insert size range that is within Z standard deviations from the mean; by default Z = 4). Plots for each alignment file are stacked and share a common x-axis that reports the chromosomal position. By convention, the sample of interest (e.g., proband or tumor) is displayed as the top track, followed by the set of related reference genomes tracks (e.g., parents and siblings, matched normal sample). Users may specify the exact order by using command line parameters to *samplot.* A visualization of all genes and exons within the locus is displayed below the alignment plots to provide context for assessing the SV's relevance to phenotypes. Rendering time depends on the number of samples and the size of the SV, but most images will require less than 5 seconds, and *samplot* rendering can be parallelizable by SV call.

**Figure 2.**
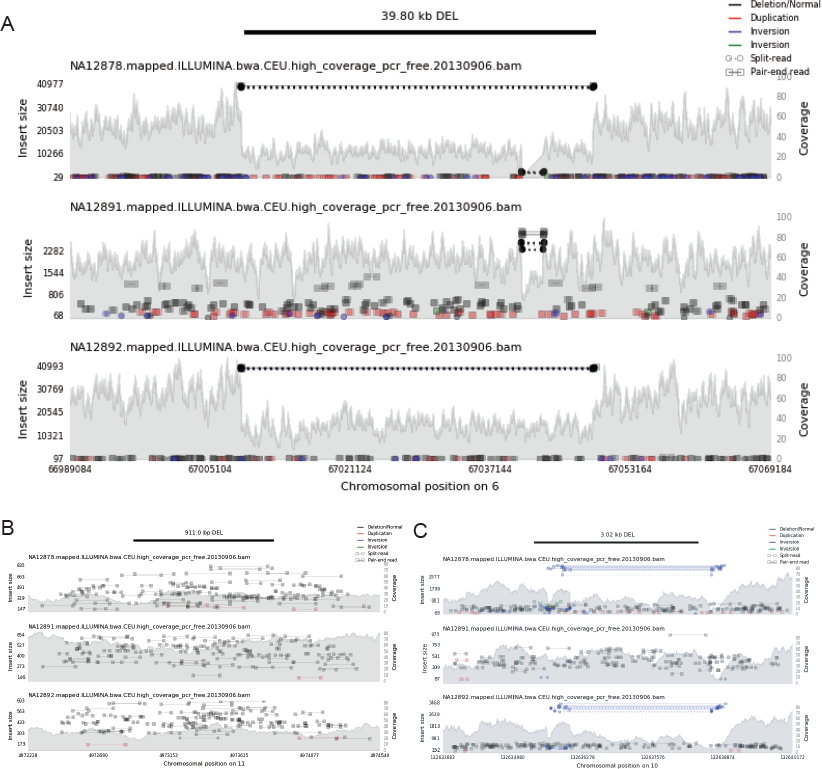
Example *samplot* images of putative deletion calls that were scored as **A)** unanimously GOOD, **B**) unanimously BAD, and **C**) ambiguous with a mix of GOOD and BAD scores. The black bar at the top of the figure indicates the genomic position of the predicted SV, and the following subfigures visualize the alignments and sequence coverage of each sample. Subplots report paired-end (square-ends connected by a solid line) and split-read (circle-ends connected by a dashed line) alignments by their genomic position (x-axis) and the distance between mapped ends (insert size, left y-axis). Colors indicate the type of event the alignment supports (black for deletion, red for duplication, and blue and green for inversion) and intensity indicates the concentration of alignments. The grey filled shapes report the sequence coverage distribution in the locus for each sample (right y-axis).

### PlotCritic

*PlotCritic* (Fig 1B) provides a simple web interface for scoring images and viewing reports that summarize the results from multiple users and SV images. *PlotCritic* is both highly scalable and easy to deploy. Images are stored on Amazon Web Services (AWS) S3 and DynamoDB tables store project configuration metadata and user responses. These AWS services allow *PlotCritic* to dynamically scale to any number of users. It also precludes the need for hosting a dedicated server, thereby facilitating deployment.

After *samplot* generates the SV images, *PlotCritic* manages their transfer to S3 and configures tables in DynamoDB based on a JSON configuration file (`config.json` file in Fig 1B). In this configuration file, one defines the curation questions posed to reviewers, as well as the allowed answers and associated keyboard bindings to allow faster responses (`curationQandA` field in Fig 1B). In turn, these dictate the text and buttons that appear on the resulting web interface. As such, it allows the interface to be easily customized to support a wide variety of curation scenarios. For example, a cancer experiment may display a tumor sample and matched normal sample and ask users if the SV appears in both samples (i.e., a germline variant) or just in the tumor sample (i.e., a somatic variant). To accomplish this, the curation question (`question` field in Fig 1B) could be “In which samples does the SV appear?”, and the answer options (`answers` field in Fig 1B) could be “TUMOR”, “BOTH”, “NORMAL”, “NEITHER”. Alternatively, in the case of a rare disease, the interface could display a proband and parents and ask if the SV is only in the proband (i.e., de novo) or if it is also in a parent (i.e., inherited). Since there is no limit to the length of a questions or number of answers options, *PlotCritic* can support more complex experimental scenarios.

Once results are collected, *PlotCritic* can generate a tab-delimited report or annotated VCF that, for each SV image, details the number of times the image was scored and the full set of answers it received. Additionally, a curation score can be calculated for each image by providing a value for each answer option and an aggregation function (e.g., mean, median, mode, standard deviation, min, max). For example, consider the cancer example from above where the values three, two, one, and zero mapped to the answers “TUMOR”, “BOTH”, “NORMAL”, and “NEITHER”, respectively. If "mode" were selected as the curation function, then the curation score would reflect the opinion of a plurality of users. The mean would reflect the consensus among all users, and the standard deviation would capture the level of disagreement about each image. While we expect mean, median, mode, standard deviation, min, and max to satisfy most use cases, users can implement custom scores by operating on the tab-delimited reported.

Each *PlotCritic* project is protected by AWS Cognito user authentication, which securely restricts access to the project website to authenticated users. A project manager is the only authorized user at startup and can authenticate other users using Cognito’s secure services. The website can be further secured using HTTPS and additional controls, such as IP restrictions, can be put in place by configuring AWS IAM access controls directly for S3 and DynamoDB.

## RESULTS

To assess *SV-plaudit’s* utility for curating SVs, nine researchers in the Quinlan laboratory at the University of Utah manually inspected and scored the 1,350 SVs (1,310 deletions, 8 duplications, 4 insertions, and 28 inversions) that the 1000 Genomes Project^1^ identified in the NA12878 genome (**Supplemental File 1**). Since we expect trio analysis to be a common use case of *SV-plaudit,* we included alignments from NA12878 and her parents (NA12891 and NA12892), and participants considered the curation questions “The SV in the top sample (NA12878) is:” and answers “GOOD”, “BAD”, or “DE NOVO”. In total, the full experiment took less than two hours with Amazon costs totaling less than $0.05. The images (**Supplemental File 2**) were generated in 3 minutes (20 threads, 2.7 seconds per image) and uploading to S3 required 5 minutes (full command list in **Supplemental File 3**). The mean time to score all images was 60.1 minutes (2.67 seconds per image) (Fig 3A, reports in **Supplemental Files 4,5**). In the scoring process, no de novo variants were identified. 40 images did not render correctly due to issues in the alignment files (e.g., coverage gaps) and were removed from the subsequent analysis (**Supplemental File 6**).

**Figure 3.**
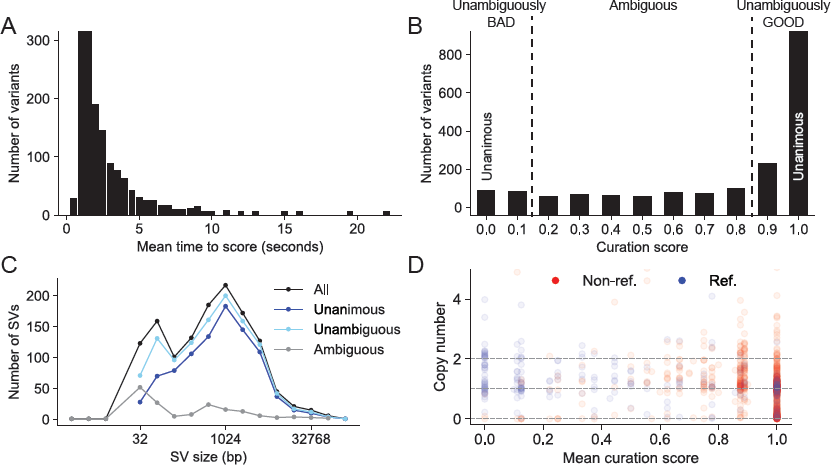
A) The distribution of the time between when an image was presented and when it was scored. B) The distribution of curation scores. C) The SV size distribution for all, unanimous (score 0 or 1), unambiguous (score <0.2 or >0.8) and ambiguous (score >=0.2 and <= 0.8) variants. D) A comparison of predictions for deletions between CNVNATOR copy number calls (y-axis), SVTYPER genotypes (color, “Ref.” is homozygous reference and “Non-ref.” is heterozygous or homozygous reference), and curation scores (x-axis). This demonstrates a general agreement between all methods. There is a concentration of reference genotypes and copy number two (no evidence for a deletion) at curation score less than 0.2. Furthermore, there is an enrichment of non-reference and copy number one or zero events at curation score greater than 0.8), false positives for CNVNATOR (copy number less than 2 at score = 0), and false negatives for SVTYPER (reference genotype at score = 1).

For this experiment, we use a curation score that mapped “GOOD” and “DE NOVO” to the value one, “BAD” to the value zero, and the mean as the aggregation function (Fig 3B). Most (70.5%) of variants were scored unanimously, with 67.1% being unanimously “GOOD” (score = 1.0, e.g., Fig 2A) and 3.4% being unanimously “BAD” (score = 0.0, e.g. Fig 2B). Since we had nine scores for each variant, we expanded our definition of “unambiguous” variants to be those with at most one dissenting vote (score <0.2 or >0.8), which accounts for 87.1% of the variants. The 12.9% of SVs that were “ambiguous” (more than one dissenting vote, 0.2<= score <=0.8) were generally small (median size of 310.5bp versus 899.5bp for all variants, Fig 3C) or contained conflicting evidence (e.g., paired-end and split-read evidence indicated an inversion and the read-depth evidence indicated a deletion, e.g., Fig 2C).

Other methods, such as SVTYPER^23^ and CNVNATOR^24^, can independently assess the validity of SV calls. SVTYPER genotypes SVs for a given sample by comparing the number of discordant paired-end alignments and split-read alignments that support the SV to the number of pairs and reads that support the reference allele. CNVNATOR uses sequence coverage to estimate copy number for the region affected by the SV. Both of these methods confirm the voting results (Fig 3D). Considering the set of “unambiguous” deletions, SVTYPER and CNVNATOR agree with the *SV-plaudit* curation score in 92.3% and 81.7% of cases, respectively. Here, agreement means that unambiguous false SVs (curation score < 0.2) have a CNVNATOR copy number near two (between 1.4 and 2.4) or an SYTYPER genotype of homozygous reference. Unambiguous true SVs (curation score > 0.8) have a CNVNATOR copy number near one or zero (less than 1.4), or an SYTYPER genotype of non-reference (heterozygous or homozygous alternate).

Despite this consistency, using either SVTYPER or CNVNATOR to validate SVs can lead to false positives or false negatives. For example, CNVNATOR reported a copy number loss for 44.2% of the deletions that were scored as unanimously BAD, and SVTYPER called 30.7% of the deletions that were unanimously GOOD as homozygous reference. Conversely, CNVNATOR had few false negatives (2.4% of unanimously GOOD deletions were called as copy neutral), and SVTYPER had few false positives (0.2% of non-reference variants were unanimously BAD).

These results demonstrate that, with *SV-plaudit,* manual curation can be a cost-effective and robust part of the SV detection process. While we anticipate that automated SV detection methods will continue to improve, due in part to the improved truth sets that *SV-plaudit* will provide, directly viewing SVs will remain an essential validation technique. By extending this validation to full call sets, *SV-plaudit* not only improves specificity but can also enhance sensitivity by allowing user to relax quality filters and rapidly screen large sets of calls. Beyond demonstrating *SV-plaudit’s* utility, our curation of SVs for NA12878 is useful as a high-quality truth set for method development and tuning. A VCF of these variants annotated with their curation score is available in **Supplementary File 5**.

## DISCUSSION

*SV-plaudit* is an efficient and scalable framework for the manual curation of large-scale SV call sets. Backed by Amazon S3 and DynamoDB, *SV-plaudit* is easy to deploy and scales to teams of any size. Each instantiation of *SV-plaudit* is completely independent and can be deployed locally for private or sensitive datasets, or be distributed publicly to maximize participation. By rapidly providing a direct view of the raw data underlying candidate SVs, *SV-plaudit* delivers the infrastructure to manually inspect full SV call sets. This functionality is vital to a wide range of WGS experiments, from method development to the interpretation of disease genomes. We are actively working on machine learning methods that will leverage the curation scores for thousands of SV prediction as training data.

*SV-plaudit* was designed to judge how well the data in an alignment file corroborates a candidate SV. The question of whether a particular SV is a false positive due to artifacts from sequencing or alignment is a broader issue that must be answered in the context of other data sources such as mappability and repeat annotations. While this second level of analysis is crucial, it is beyond the scope of this paper, and we argue this analysis be performed only for those SVs that are fully supported by the alignment data. While *SV-plaudit* combines *samplot* and *PlotCritic* to enable the curation of structural variant images, we emphasize that the *PlotCritic* framework can be used to score images of any type. Therefore, we anticipate that this framework will facilitate "crowd-sourced" curation of many other biological images.

